# Hospitalized premature infants are colonized by related bacterial strains with distinct proteomic profiles

**DOI:** 10.1101/217950

**Authors:** Christopher. T. Brown, Weili Xiong, Matthew R. Olm, Brian C. Thomas, Robyn Baker, Brian Firek, Michael J. Morowitz, Robert L. Hettich, Jillian F. Banfield

**Affiliations:** Department of Plant and Microbial Biology, University of California, Berkeley, CA, USA; Biosciences Division, Oak Ridge National Laboratory, Oak Ridge, Tennessee, USA; Department of Earth and Planetary Science, University of California, Berkeley, CA, USA; Division of Newborn Medicine, Children’s Hospital of Pittsburgh and Magee-Womens Hospital of UPMC, Pittsburgh,United States; Department of Surgery, University of Pittsburgh School of Medicine, Pittsburgh, PA, USA; Division of Pediatric General and Thoracic Surgery, Children’s Hospital of Pittsburgh of UPMC,Pittsburgh, PA, USA; Chemical Science Division, Oak Ridge National Laboratory, Oak Ridge, Tennessee, USA; Department of Environmental Science, Policy, and Management, University of California, Berkeley, CA, USA; Earth Sciences Division, Lawrence Berkeley National Laboratory, Berkeley, CA, USA

**Keywords:** Metagenomics, Metaproteomics, Microbial Genomics, iRep, Human Microbiome, Necrotizing Enterocolitis, Neonates, Microbial Colonization

## Abstract

During the first weeks of life, microbial colonization of the gut impacts human immune system maturation and other developmental processes. In premature infants, aberrant colonization has been implicated in the onset of necrotizing enterocolitis (NEC), a life-threatening intestinal disease. To study the premature infant gut colonization process, genome-resolved metagenomics was conducted on 343 fecal samples collected during the first three months of life from 35 premature infants housed in a neonatal intensive care unit, 14 of which developed NEC, and metaproteomic measurements were made on 87 samples. Microbial community composition and proteomic profiles remained relatively stable on the time scale of a week, but the proteome was more variable. Although genetically similar organisms colonized many infants, most infants were colonized by distinct strains with metabolic profiles that could be distinguished using metaproteomics. Microbiome composition correlated with infant, antibiotics administration, and NEC diagnosis. Communities were found to cluster into seven primary types, and community type switched within infants, sometimes multiple times. Interestingly, some communities sampled from the same infant at subsequent time points clustered with those of other infants. In some cases, switches preceded onset of NEC; however, no species or community type could account for NEC across the majority of infants. In addition to a correlation of protein abundances with organism replication rates, we found that organism proteomes correlated with overall community composition. Thus, this genome-resolved proteomics study demonstrates that the contributions of individual organisms to microbiome development depend on microbial community context.

**Importance**

Humans are colonized by microbes at birth, a process that is important to health and development. However, much remains to be known about the fine-scale microbial dynamics that occur during the colonization period. We conducted a genome-resolved study of microbial community composition, replication rates, and proteomes during the first three months of life of both healthy and sick premature infants. Infants were found to be colonized by similar microbes, but each underwent a distinct colonization trajectory.

Interestingly, related microbes colonizing different infants were found to have distinct proteomes, indicating that microbiome function is not only driven by which organisms are present, but also largely depends on microbial responses to the unique set of physiological conditions in the infant gut.

## Introduction

Infants have high levels of between-individual variation in microbiome composition compared with adult humans (1, 2). Variation in the infant microbiome exists at both the species and strain level (3, 4). During the first one to two years of life, the gut microbiome of infants begins to converge upon an adult-like state (2, 5). However, aberrations in this process may contribute to diseases such as type 1 and 2 diabetes, irritable bowel disease, and necrotizing enterocolitis (NEC) in premature infants (6-11). Because establishment of the microbiome is a key driver of immune system development, changes in the process of colonization may have life-long implications, even if they do not result in drastically different microbiome composition later in life (12, 13).

Infants born prematurely have lower diversity microbial communities compared with full term infants, and are susceptible to life-threatening diseases such as NEC (4, 14-17). While it has long been thought that bacterial infection may contribute to NEC pathogenesis, strain-resolved microbial community analysis has not identified a single pathogen that is responsible for the disease (3). However, it is still likely that microbial communities play an important role, with the context-dependent metabolism of specific strains potentially critical to infant health and disease. Recent studies have applied proteomics and metabolomics to premature infant gut microbiomes to measure functional profiles in healthy premature infants and those that went on to develop NEC (18, 19).

These studies reported temporal variation in the infant proteome and identified metabolites associated with NEC. However, further study is required to better understand the range of functional and developmental patterns during the microbial colonization process.

To investigate microbial community assembly, and how microbes modulate their metabolism and replication rate during colonization, we conducted a combined metagenomics and metaproteomics study of the microbiome of both healthy premature infants and infants that went on to develop NEC. Microbiome samples were collected during the first three months of life with the goal of measuring the physiological changes of dominant and ubiquitous bacterial species. Genomes assembled from metagenomes enabled analysis of microbial community membership, and tracking of both community composition and replication rates over time. The availability of genome sequences made it possible to map protein abundance measurements to bacterial species and strains.

Microbial communities were clustered into distinct types in order to provide context for proteomics analyses. Statistical analyses showed that, while species and strain-specific proteomic profiles correlated with overall community composition, the proteomes of members of the same species and strain were largely infant-specific. These analyses also show that bacterial proteome features are correlated with infant development, health status, and antibiotics administration.

## Results

### Metagenome sequencing and genome binning

In order to study the developing gut microbiome, stool samples were collected during the first three months of life for 35 infants born prematurely and housed in the neonatal intensive care unit at Magee-Womens Hospital at the University of Pittsburgh Medical Center. Two of the infants in the study cohort developed sepsis (N1_017 and N1_019) and 14 infants developed necrotizing enterocolitis (NEC; **Table 1**). To study the gut microbiome, we analyzed 1,149 Gbp of DNA sequences generated by our laboratory (3, 4, 20). These sequences were from 343 metagenomes (average of 3.3 Gbp sequencing per sample; **Supplemental Figure 1 and Supplemental File 1a**). Metagenomes were assembled into 6.79 Gbp of scaffolds ≥1 Kbp that represented 92% of all sequenced DNA.

**Table 1.** | Infant medical information.

| infant | campaign | sex | delivery | mult. gest. | gestational<br>age<br>(weeks) | birth<br>weight (g) | feeding | condition | NEC<br>diagnosis<br>(DOL) |
| --- | --- | --- | --- | --- | --- | --- | --- | --- | --- |
| N1_003 | NIH1 | F | C-section | Single | 26 | 822 | Breast | control | n/a |
| N1_004 | NIH1 | F | C-section | N1_005 | 32 | 1450 | Formula | control | n/a |
| N1_008 | NIH1 | F | Vaginal | Single | 32 | 1230 | Formula | NEC | 9 |
| N1_009 | NIH1 | M | C-section | Single | 29 | 1820 | Combination | control | n/a |
| N1_011 | NIH1 | M | C-section | N1_012 | 26 | 523 | Combination | NEC | 34, 62 |
| N1_014 | NIH1 | M | Vaginal | Single | 32 | 2035 | Combination | control | n/a |
| N1_017 | NIH1 | F | Vaginal | Single | 26 | 748 | Combination | NEC | 11 |
| N1_018 | NIH1 | M | C-section | Single | 29 | 1133 | Combination | control | n/a |
| N1_019 | NIH1 | F | C-section | N1_020, N1_021 | 24 | 731 | Combination | control | n/a |
| N1_021 | NIH1 | F | C-section | N1_019, N1_020 | 24 | 697 | Breast | NEC | 32 |
| N1_023 | NIH1 | F | Vaginal | Single | 27 | 875 | Breast | control | n/a |
| N2_031 | NIH2 | M | C-section | Single | 26 | 773 | Formula | control | n/a |
| N2_035 | NIH2 | M | Vaginal | Single | 25 | 795 | Breast | control | n/a |
| N2_038 | NIH2 | F | C-section | N2_039 | 30 | 1381 | Combination | control | n/a |
| N2_039 | NIH2 | F | C-section | N2_038 | 30 | 1470 | Combination | NEC | 24 |
| N2_060 | NIH2 | M | C-section | Single | 30 | 1878 | Combination | control | n/a |
| N2_061 | NIH2 | M | Vaginal | Single | 28 | 1184 | Combination | NEC | 9, 34 |
| N2_064 | NIH2 | M | Vaginal | Single | 28 | 1100 | Combination | control | n/a |
| N2_065 | NIH2 | F | Vaginal | Single | 25 | 841 | Combination | control | n/a |
| N2_066 | NIH2 | F | Vaginal | Single | 28 | 1028 | Breast | control | n/a |
| N2_069 | NIH2 | M | C-section | N2_070 | 26 | 637 | Breast | NEC | 32 |
| N2_070 | NIH2 | F | C-section | N2_069 | 26 | 633 | Combination | control | n/a |
| N2_071 | NIH2 | M | C-section | Single | 25 | 754 | Combination | NEC | 31 |
| N2_088 | NIH2 | F | C-section | N2_089 | 28 | 1057 | Formula | control | n/a |
| N2_093 | NIH2 | M | C-section | Single | 26 | 924 | Breast | NEC | 12 |
| N3_172 | NIH3 | M | C-section | Single | 28 | 1250 | Breast | NEC | 37, 54 |
| N3_173 | NIH3 | M | C-section | Single | 29 | 1530 | Breast | NEC | 25 |
| N3_174 | NIH3 | F | C-section | Single | 30 | 980 | Breast | control | n/a |
| N3_175 | NIH3 | M | Vaginal | Single | 29 | 1480 | Combination | control | n/a |
| N3_176 | NIH3 | M | C-section | Single | 28 | 990 | Combination | control | n/a |
| N3_177 | NIH3 | F | Vaginal | Single | 28 | 900 | Combination | control | n/a |
| N3_178 | NIH3 | M | Vaginal | Single | 32 | 2050 | Combination | NEC | 16 |
| N3_182 | NIH3 | M | C-section | Single | 39 | 3010 | Combination | NEC | 6 |
| N3_183 | NIH3 | M | Vaginal | Single | 32 | 2410 | Combination | NEC | 11 |
| S2_010 | NIH3 | M | C-section | Single | 32 | 1810 | Combination | control | n/a |

**Figure 1.**
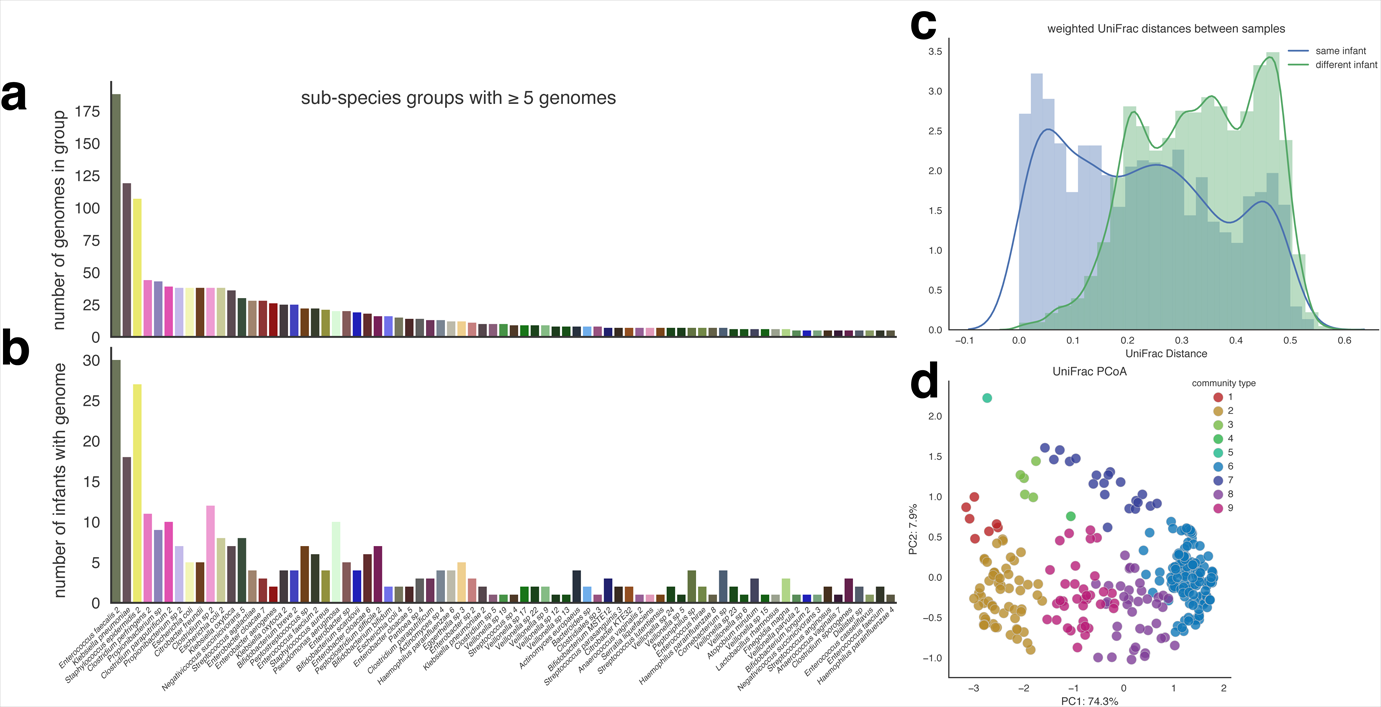
| Premature infant gut microbial communities associated into seven primary types. Genomes reconstructed from metagenomes were clustered into sub-species groups based on sharing 98% average nucleotide identity (ANI). **a**, The number of genomes assigned to each group and **b**, the number of infants with a reconstructed genome from the group. Shown are groups comprised of five or more genomes. **c**, Pairwise weighted UniFrac distances calculated between all microbiome samples based on genome sequence ANI and abundance. **d**, PCoA clustering of samples based on weighted UniFrac distances. Samples are colored based on community type assignment.

Scaffolds assembled from metagenomes were grouped into 3,643 bins, 1,457 of which were ≥50% complete with ≤5% contamination; **Supplemental Figure 2**, **Supplemental File 2**). These genomes were assigned to 270 groups approximating different bacterial sub-species based on sharing ≥98% average nucleotide identity (ANI) (**Supplemental File 1b**). These genomes account for 91% of the total sequencing.

**Figure 2.**
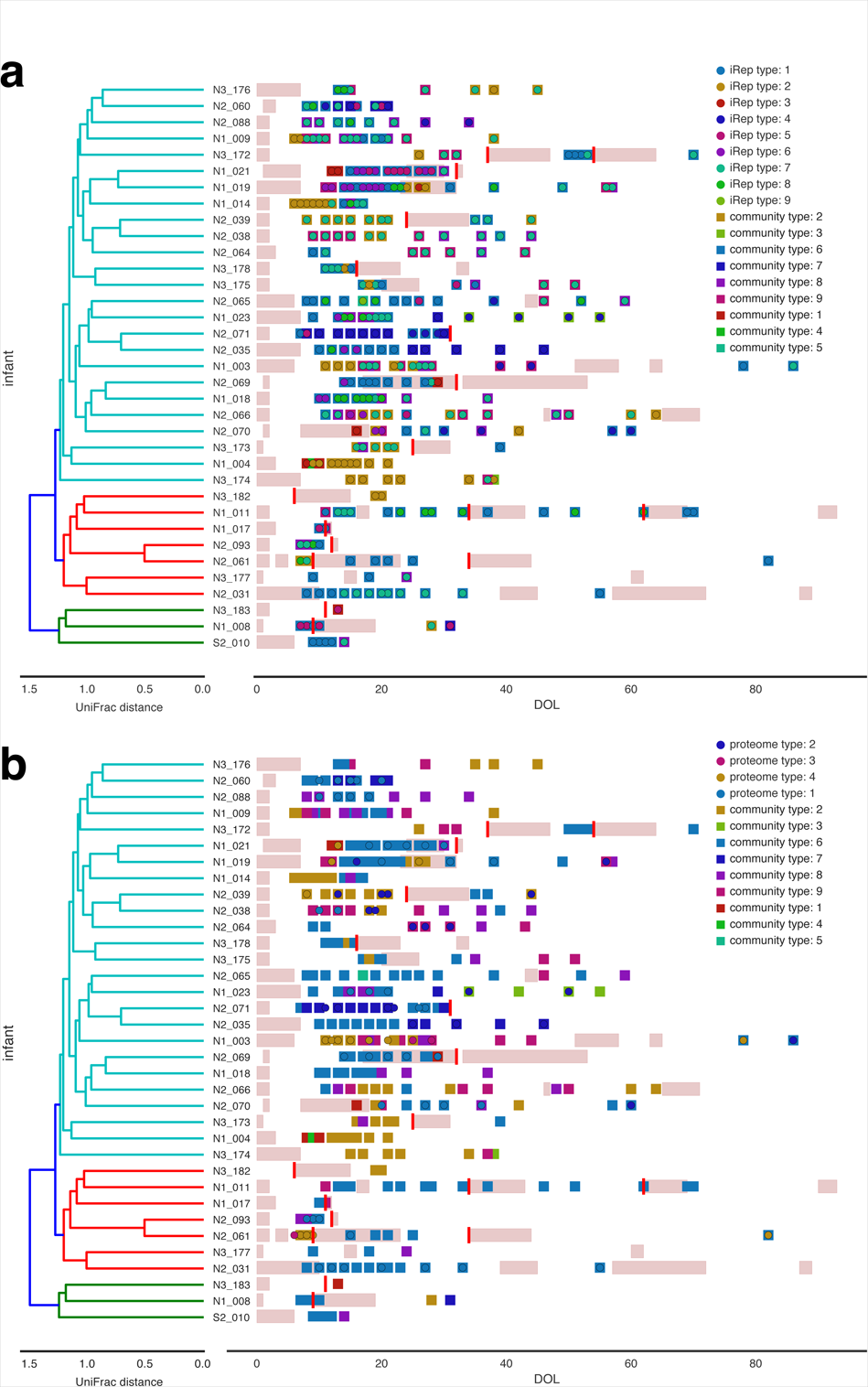
| Microbial colonization patterns for preterm infants. Samples were clustered into types based on microbial community composition (“community type”), bacterial iRep profiles (“iRep type”), and overall bacterial proteome composition (“proteome type”). Microbial community type is shown along with iRep **(a)** and proteome **(b)** types. Infants are arranged based on hierarchical clustering of unweighted UniFrac distances calculated based on the set of genomes recovered from each infant (**Supplemental Figure 3**). Antibiotics administration is indicated with pink bars and NEC diagnoses with red bars. DOL stands for day of life.

Genomes suitable for iRep replication rate analysis (≥75% complete with ≤175 fragments/Mbp and ≤5% contamination) were available for 193 genome clusters (21). ***Protein quantification by metaproteomics***

Across all metagenomes, 5,233,047 proteins were predicted, 897,520 of which were from a non-redundant set of representative genomes clustered at 98% ANI. Proteins clustered into 121,746 putative families (**Supplemental File 3**). Metaproteomics measurements were conducted on 87 metagenome-matched samples that spanned 16 infants, six of which developed NEC and one of which was diagnosed with sepsis (N1_019; **Supplemental Figure 1**). Conducting metagenomics and metaproteomics on the same samples was critical for obtaining an appropriate database for matching peptides to proteins. On average, 71,676 unique bacterial spectral counts were detected per sample, with an average of 33% of predicted bacterial proteins identified (**Supplemental Figure 1**, **Supplemental File 1b, and Supplemental File 4**).

### Premature infants were colonized by genetically similar organisms, and microbial communities clustered into seven primary types

The majority of infants were colonized by *Enterococcus faecalis, Klebsiella pneumoniae*, and *Staphylococcus epidermidis* (**Figure 1a,b**). Overall, infants that developed NEC were colonized by organisms genetically similar to those colonizing other infants, and most genotypes were seen in only one infant. No individual species was strongly associated with NEC (**Supplemental Figure 3**).

**Figure 3.**
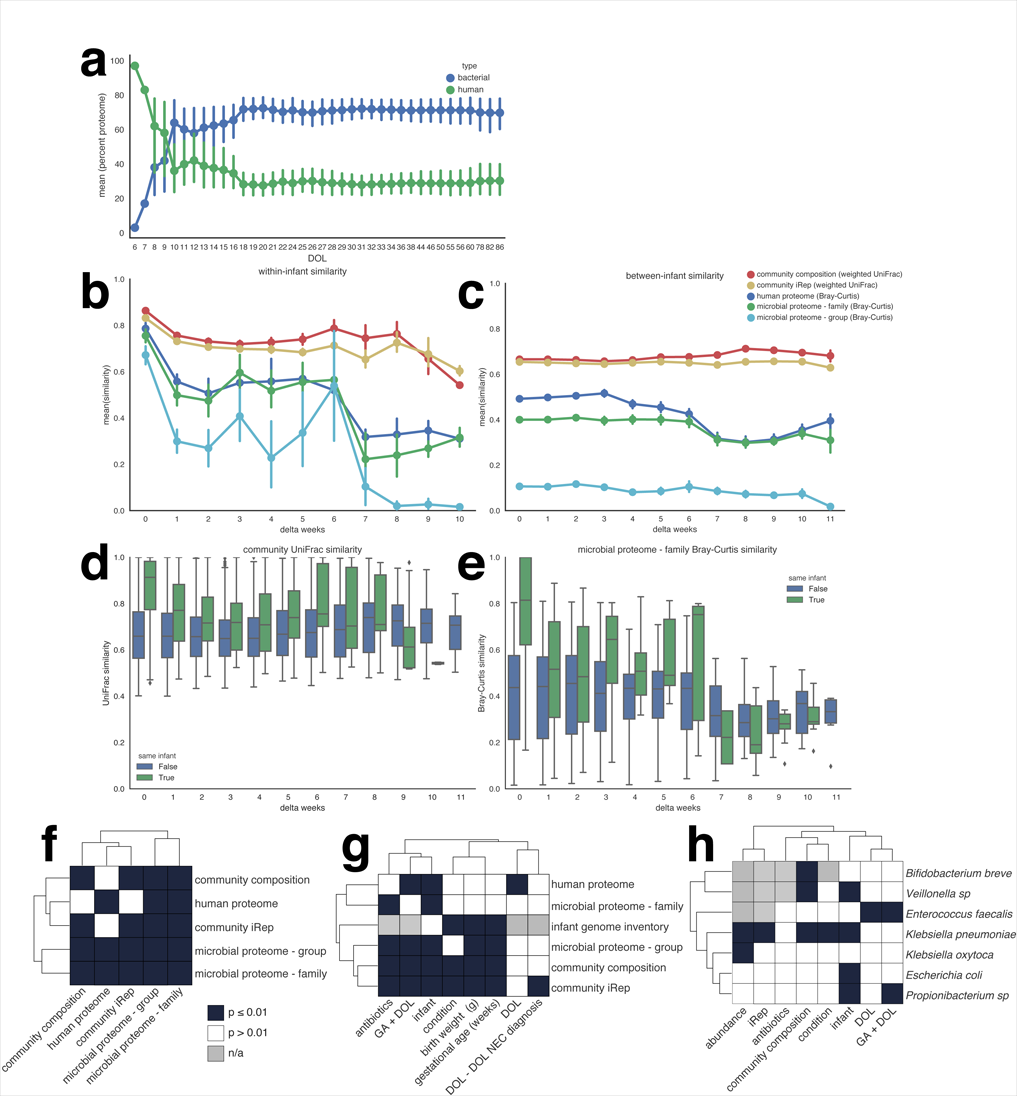
| Microbiome stability and correlations. **a**, The relative contribution of human and bacterial proteins to overall proteome composition during development of the premature infant gut. **b**, Similarity measurements for microbiomes sampled either from the same infant or **c**, from different infants. Comparison of similarity measurements calculated between samples collected either form the same or different infants based either on weighted microbial community UniFrac (**d**), or weighted microbial proteome BrayCurtis (**e**) measurements. Human proteome and microbial community correlations calculated between one another (**f**), with infant metadata (**g**), and determined based on microbial species (**h**). Shown are PERMANOVA or Mantel test p-values (**f-h**). Microbial proteome “family” refers to protein family analysis, and “group” refers to analysis of proteins clustered at 97% amino acid identity.

The premature infant microbiome was found to be highly variable. In some cases, samples collected from an infant at subsequent time points were as different from earlier samples as those collected from other infants (**Figure 1c**). Communities were clustered based on species membership and abundance in order to identify microbial consortia common during the colonization process. In order to account for both genomic differences and organism abundance, clustering was conducted based on weighted UniFrac distances, where the tree used for calculating UniFrac was constructed using genome ANI. Nine distinct community types were identified, seven of which were comprised of samples collected from multiple infants and were thus considered primary types (**Figure 1d, Supplemental Figure 4, and Supplemental Figure 5**). Each community type is characterized by the dominance of different community members (**Supplemental Figure 6**). Microbiomes from different infants clustered into the same community type, and the microbiome of individual infants was found to switch types, sometimes multiple times, during the colonization process (**Figure 2**). Although infants shared community types, overall colonization patterns were not replicated across infants. Microbiomes associated with infants that did and did not go on to develop NEC were often classified in the same community type. In some cases, switches preceded onset of NEC, but no type or switch could explain all cases of NEC.

**Figure 4.**
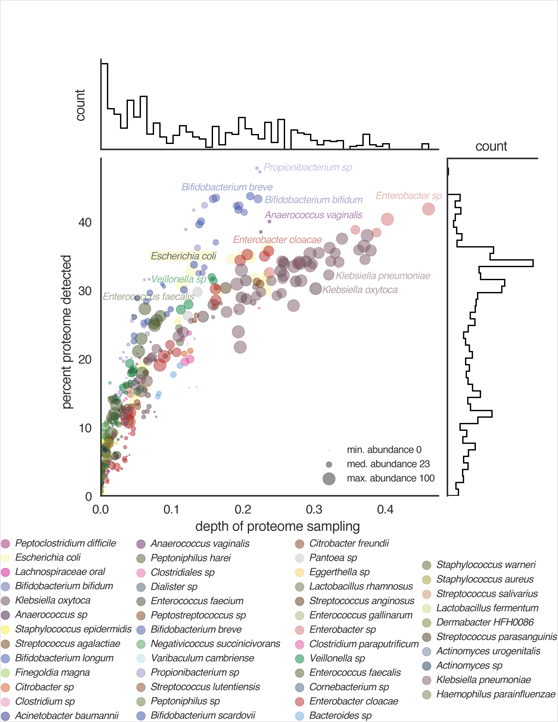
| Proteome detection for species colonizing premature infants. Depth of proteome sampling for organisms in each sample is compared against the percent of predicted proteins that could be detected. Data point sizes and histograms are scaled based on organism abundance as determined by metagenome sequencing.

**Figure 5.**
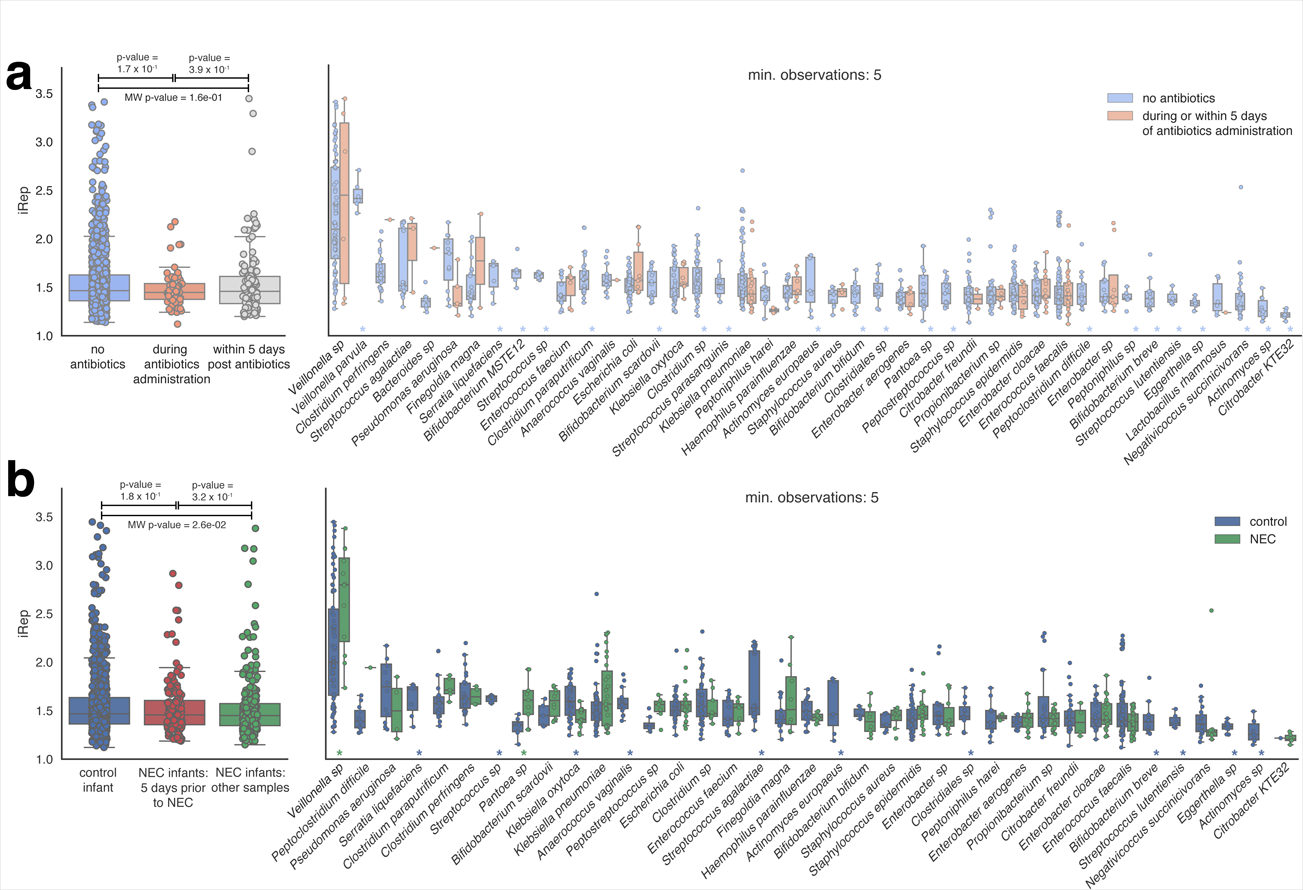
| Replication rates for bacteria colonizing premature infants. **a**, Replication rates for bacteria sampled during periods with or without antibiotics administration and **b**, associated with infants that did and did not go on to develop NEC. Statistically significant differences between replication rates observed for individual species under different conditions are indicated with an asterisk (MW p-value ≤0.01). Shown are all species with at least five observations.

**Figure 6.**
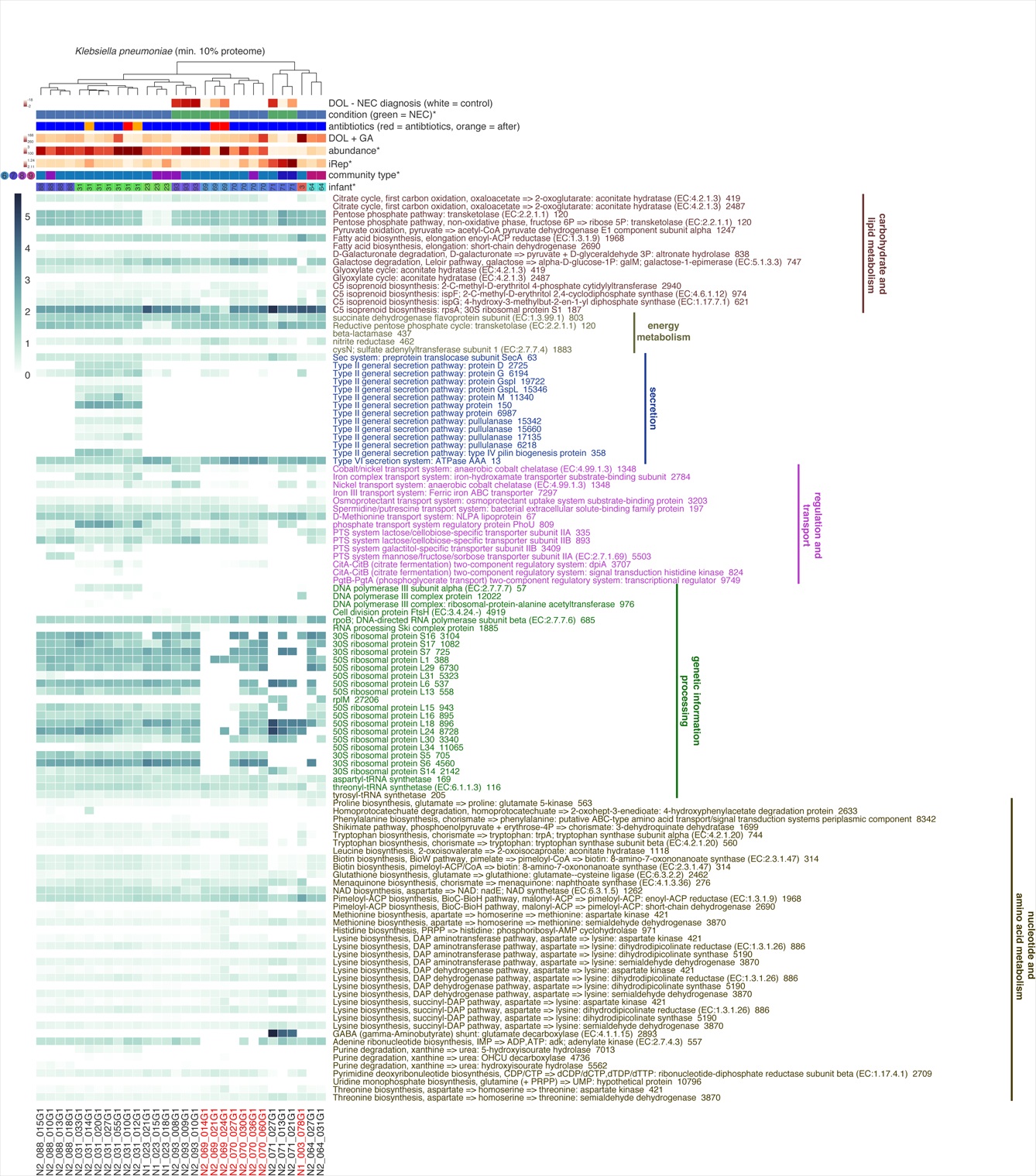
| *Klebsiella pneumonia* proteins with infant-specific abundance profiles. Hierarchical clustering was conducted on all *K. pneumonia* protein families, showing that strains colonizing different infants have distinct proteomic profiles. Infant and species metadata are shown for each sample. Metadata significantly correlated with the *K. pneumonia* proteome are indicated with an asterisk (PERMANOVA or Mantel test p-value ≤0.01). Protein families that correlated with at least one infant are shown in the heatmap (edgeR q-value ≤0.01). Samples colonized by the same *K. pneumonia* strain are shown with red text.

### Microbial replication rates and proteins

iRep is a newly-developed method that enables measurement of bacterial replication rates based on metagenome sequencing data when high-quality draft genome sequences are available (21). We applied the iRep method using genomes recovered from metagenomes sequenced for each infant in the study, and quantified 1,328 iRep replication rates from 330 samples. Sample clustering was conducted based on community iRep profiles, identifying nine distinct iRep types that were correlated with community type (Mantel test p-value = 1 x 10^-3^, **Figure 2a**). Likewise, analysis of protein family abundance clustered samples into four distinct proteome types, which also correlated with community type (Mantel test p-value = 1 x 10^-3^, **Figure 2b**). Interestingly, there are several cases in which iRep and/or proteome type switched when community type was constant, or when community type switched but iRep and/or proteome type remained constant.

### Microbiome development

Peptide spectral counts were matched to infant-specific databases containing both human and microbial proteins. This allowed for the relative proportions of human and microbial proteins to be determined for each time point. Samples are dominated by human proteins during the first 10 days of life (DOL), and then microbial proteins become dominant around DOL 18. Ratios of human versus bacterial protein abundances show that the premature infant gut microbiome is established over a period of approximately two weeks (**Figure 3a**).

The presence of multiple data types (microbial community abundance and iRep, microbial community proteome composition, and human proteome composition) enabled tracking of various aspects of human and microbiome development during the first months of life (**Figure 3b,c**). All measurements from an infant were stable within the time scale of a week, but diverged over time. Interestingly, communities from different infants neither converged nor diverged over time in terms of similarity based on three of these five metrics. However, we observed that human proteome measurements and microbial protein family abundances from different infants became increasingly different when samples with time separations of greater than three weeks were compared. Overall, the microbial proteome was more variable (higher variance) than community composition (**Figure 3d,e**). After approximately two weeks, both microbial community abundance and proteome measurements collected from the same infant became as different from each other as samples collected from other infants.

The majority of human and microbiome features recorded in our analyses were correlated with one another (**Figure 3f**). However, an exception is that microbial community abundance and iRep were not correlated with human proteome composition (Mantel test p-value >0.01). This is interesting in that it shows that there is no strong connection between the overall human proteome and either the composition or replication activity of the microbiome.

As shown in **Figure 3g**, microbial features were also correlated with a variety of infant factors, including infant health and development (gestational age and weight), as well as antibiotics administration (Mantel or permutational analysis of variance, PERMANOVA, p-value ≤0.01). Notably, whether or not an infant developed NEC (condition) correlated with several microbiome factors (infant genome inventory, and both community composition and iRep), but not with proteome measurements. However, these correlations were in part driven by antibiotics, as only iRep was correlated with infant condition when excluding samples collected during or within five days of antibiotics administration. Regardless of the influence of antibiotics on the microbiome, microbial responses to treatment likely impact infant health.

### Different species expressed varying amount of their proteome in the infant gut

Microbes present in the gut environment are not expected to express their complete complement of proteins at all times. In order to investigate the extent of proteome expression for different bacteria, we compared depth of proteome sampling for each organism to the percent of the predicted proteome that could be detected (**Figure 4**). The median proteome detection across all samples was 11%, but this was largely due to low sampling depth. Higher depth of proteome sampling corresponded with detection of a larger fraction of the predicted proteins. The median percent of the proteome detected for organisms with the best detection in each sample was 31% (max. 48%). For several frequently detected colonists, including *Klebsiella pneumoniae, Klebsiella oxytoca*, and members of the genus *Enterobacter*, maximum proteome expression was ∼50%. However, *Propionibacterium sp., Anaerococcus vaginalis*, and members of the genus *Bifidobacterium* expressed a greater proportion of their encoded genes than other organisms. We infer that these bacteria may be specifically adapted to environments and resource availability within the infant gut, whereas other bacteria may maintain capacities that enable adaption to other environments.

### Members of the same bacterial species replicated at different rates during colonization

Across all infants, *Streptococcus agalactiae, Pseudomonas aeruginosa, Klebsiella pneumoniae*, and members of the genera *Veillonella* and *Clostridium* exhibited some of the highest replication rates (**Supplemental File 1c**). iRep values for organisms sampled in this cohort during or immediately after antibiotics administration were not significantly different from those at other time points (**Figure 5a**). This indicates that populations present after antibiotics administration are both resistant to antibiotics and are continuing to replicate. Members of several species were replicating quickly during or immediately following antibiotic treatment (*Veillonella sp., Streptococcus agalactiae, Finegoldia magna*, and others). However, we did not detect overall higher iRep values following antibiotics administration, although this was reported previously (21). Most species were found only to be replicating in the absence of antibiotics, consistent with their susceptibility to the treatment.

### Species-specific proteomic profiles are associated with infant and microbiome features

Relative protein abundance levels were determined for each genome and tracked across samples. This identified population-specific proteome profiles and enabled us to test for correlations with various human and microbial properties (**Figure 3h**, **Supplemental Figure 1**, **and Supplemental File 1d**). *Veillonella spp., Klebsiella pneumoniae, Escherichia coli*, and *Propionibacterium sp.* all had infant-specific profiles (PERMANOVA p-value ≤0.01), indicating that although similar organisms are colonizing different infants, each population is expressing a different complement of proteins. *K. pneumoniae* and *Veillonella spp.* proteomes also correlated with community type, as did the *Bifidobacterium breve* proteome (Mantel test p-value ≤0.01), showing that populations respond to overall microbial community context. Interestingly, both *Enterococcus faecalis* and *Propionibacterium sp*. exhibited proteomes that were also correlated with infant development, and the *K. pneumoniae* proteome correlated with both iRep and infant health. Although overall microbial proteome correlated with antibiotics administration, species-specific proteome profiles did not; however, this may be due to a lack of available data for the same species in multiple samples with and without antibiotics.

Because of the existence of 35 samples in which ≥10% of the *K. pneumoniae* proteome could be detected (max. = 38%, median = 25%), correlations between individual protein abundances and iRep could be determined. Amongst proteins positively correlated with iRep were a transcriptional regulator (LysR), proteins involved in cell wall biogenesis, and ribosomal proteins (Pearson ≥0.5, q-value ≤0.01, observed in ≥15 samples; **Supplemental File 1e**).

### Infants were colonized by different strains with distinct proteomes

The finding that *K. pneumoniae, E. coli, Propionibacterium sp.*, and *Veillonella spp.* have infant-specific proteomes raised the question of whether or not each infant was being colonized by different strains. Genome sequences ≥50% complete with ≤5% contamination that were assembled for each species from each infant were compared with one another, and hierarchical clustering conducted on pairwise ANI values was used to delineate strains (**Supplemental Figure 7**). Clustering showed that in most cases each infant was indeed colonized by distinct strains, which proteomics analysis showed are functionally distinct. However, there were a few notable exceptions. Twin infants N2_069 and N2_070, as well as infant N1_003 were all colonized by the same strain of *K. pneumoniae*. The proteomic profiles for the strains colonizing N2_069 and N2_070 were more similar to one another than they were to profiles recovered from other strains; however, they were still distinguishable (**Figure 6**). Likewise, the same strain of *Propionibacterium sp.* colonized twin infants N2_038 and N2_039. As with shared strains of *K. pneumoniae*, their functional profiles clustered together but were still distinguishable from one another (**Supplemental Figure 8**).

**Figure 7.**
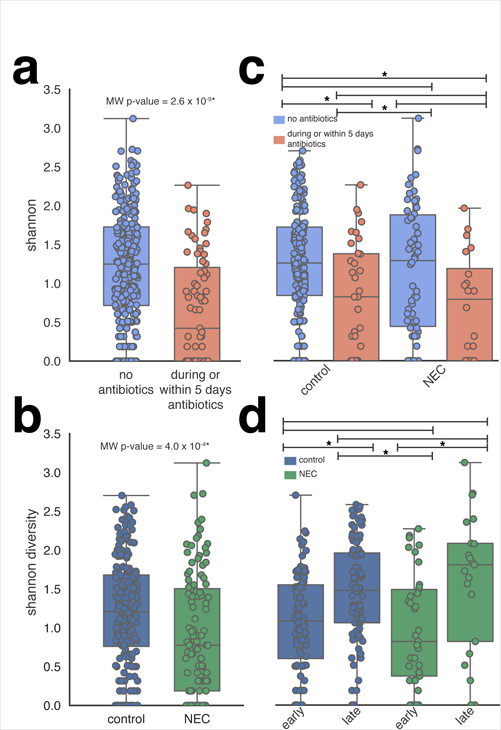
| Microbial community diversity. Shannon diversity measurements for microbial communities associated with infants during periods with or without antibiotics administration (**a**), and between infants that did and did not go on to develop NEC (**b-d**). Significant differences are indicated with an asterisk (MW p-value ≤0.01). “Early” samples were collected prior to GA + DOL 220. Samples collected after NEC diagnosis were excluded from **c** and **d**.

Analysis showed that few proteins were responsible for distinguishing proteomes of the same bacterial types in different infants (**Figure 6, Supplemental Figure 8, and Supplemental File 1d**). Common amongst these were proteins involved in nucleotide, amino acid, carbohydrate and lipid metabolism. Also notable were several proteins produced by *K. pneumoniae* involved in central carbohydrate metabolism, and both galactose degradation and D-galacturonate degradation, indicating different carbon preferences for strains colonizing different infants (**Figure 6**). Proteins involved in bacterial secretion were differentially abundant between *K. pneumonia* colonizing different infants, indicating variations in secretion potential that could affect human-microbe interactions. Relatedly, the abundance of proteins involved in transport of metals, ions, citrate, and several sugars also differed between infants.

### Low microbiome diversity was associated with both antibiotics administration and NEC

Microbiome diversity was lower during or within five days of antibiotics administration compared with other time points (Mann-Whitney U test, MW, p-value = 2.6 x 10^-9^; **Figure 7a**), and the microbiome of infants that developed NEC was typically less diverse compared with healthy infants (MW p-value = 4 x 10^-4^; **Figure 7b**). However, the difference in diversity between healthy and NEC infants was driven by the fact that NEC infants more frequently receive antibiotics (**Figure 2**). When comparing within either periods with or without antibiotics, microbiome diversity for healthy and NEC infants (pre-NEC diagnosis) was indistinguishable (**Figure 7c**). When excluding antibiotics samples, both groups of infants had higher diversity microbial communities later in development (post GA + DOL 220; **Figure 7d**).

### Microbial community composition was correlated with infant health

Premature infants that developed NEC had different microbial community abundance profiles (PERMANOVA p-value = 3 x 10^-3^; **Supplemental Figure 4g**). Interestingly, there were a variety of species detected in healthy infants, but never detected in those that developed NEC; however, the opposite was not true. It should be noted that species unique to healthy infants were not consistently detected. No species identified five days prior to NEC diagnosis showed a significant difference in abundance, or was unique to NEC infants.

Overall community composition was also correlated with each infant, antibiotics administration, birth weight, gestational age, and gestational age corrected day of life (GA + DOL; PERMANOVA or Mantel test p-value ≤0.01; **Supplemental Figure 4**). Several species were more abundant members of communities associated with infants that developed NEC: *Enterobacter sp., Propionibacterium sp.*, and *Peptostreptococcus sp.* (edgeR q-value ≤0.01 after excluding samples collected within five days of antibiotics administration; **Supplemental File 1f**). *Vellonella sp.* replicated faster in NEC infants, while several groups of organisms were replicating faster in control infants, including members of the genera *Anaerococcus, Klebsiella, Actinomyces, Eggerthella, Streptococcus, Clostridiales*, and *Bifidobacterium* (MW p-value ≤0.01 after excluding samples collected within five days of antibiotics administration; **Supplemental File 1c**). Several different species were active in control infants, but were not detected in infants that went on to develop NEC. However, combined iRep values collected from infants that did and did not go on to develop NEC were not statistically different, even when considering only samples collected within the five days prior to NEC diagnosis (**Figure 5b**).

### Microbial proteins associated with proteome type, antibiotics administration, and NEC

As described above, we used protein abundance patterns to cluster microbial community proteomes into functionally distinct proteome types. Statistical analysis identified 3,085 differentially abundant proteins distinguish proteome types (edgeR q-value ≤ 0.01; **Supplemental File 1g**). Of these, 461 were found to distinguish only one proteome type from all others. Notable amongst all of these proteins were those involved in central carbohydrate metabolism and energy metabolism (**Supplemental Figure 9**). Proteome types differ in terms of the amount and variety of carbon degradation enzymes, as well as the propensity for aerobic versus anaerobic respiration (based on the abundance of oxidases and reductases).

Samples collected during antibiotics treatment were enriched in 56 different proteins (identified in more than one treated infant, edgeR q-value ≤0.01; **Supplemental File 1g**). Amongst these proteins were those involved in secretion, transcription, and DNA degradation. Along with iRep results, the findings indicate that a subset of organisms remain active in the presence of antibiotics.

Although overall community proteome abundance profiles were not correlated with NEC, microbial proteins from 160 different protein families, many with no known function, were more abundant in samples from infants that went on to develop NEC (identified in more than one NEC infant, edgeR q-value ≤0.01; **Supplemental File 1g**). Annotated proteins were dominantly involved in transport of ions, metals, and other substrates, iron acquisition, and both motility and chemotaxis. Among proteins responsible for iron scavenging was subunit E of enterobactin synthase, a high-affinity siderophore involved in iron acquisition, which is often used by pathogenic organisms. Also more abundant was outer membrane receptor FepA, which is involved in transporting iron bound by extracellular enterobactin. Subunit F of enterobactin synthase was also identified in NEC infants, as were an iron-enterobactin ABC transporter substrate-binding protein, and an enterobactin esterase. The abundance of this protein suggests a possible role for iron acquisition by organisms that may contribute to disease onset. Interestingly, 21 *K. pneumoniae* proteins were correlated with NEC, including a ferrous iron transporter (family 2834) that was 3.9-fold more abundant in two infants that developed NEC. The abundance of this protein was also correlated with infant, proteome type, community type, and antibiotics administration.

## Discussion

Most studies to date have focused on the composition of the gut microbiome, typically at the low-resolution afforded by 16S rRNA gene amplicon methods. We used genome-resolved time-series metagenomics in conjunction with iRep replication rate and metaproteomics measurements to obtain a more comprehensive view of the colonization process. The dataset included information about the gut colonization trajectories of both healthy infants and infants that went on to develop NEC, enabling exploration of microbiome variability, at both the community composition and organism functional levels.

Microbial communities were classified into types based on the mixture of organisms present. Interestingly, most types occurred in multiple infants, a result that indicates the tendency of gut colonizing bacteria to form networks of interaction, possibly based on metabolic complementarity. An important factor determining the community type present may be the specific organisms that are introduced, and the extent to which they are able to colonize. Other factors that may dictate the community type include human genetic selection, diet, and antibiotics administration. Within a single infant, community types often switched several times over the observation period. Given the lack of evidence for consistent transitions from one type to another across multiple infants, the high degree of variation in iRep replication rates observed for members of the same species, and a lack of convergence of communities in different infants, we conclude that colonization is a chaotic process.

Overall microbial physiology, as measured by whole proteome abundance patterns, was more dynamic than community composition. Thus, metagenomics-enabled proteomic analyses indicate functional flexibility that does not depend on addition or loss of organisms. Shifts in the importance of specific pathways or metabolisms with environmental conditions would not be apparent in studies that only use organism identification or metabolic potential predictions.

It is possible that onset of NEC is due to fast growth rates of potential pathogens within communities that are imbalanced due to low species richness, ultimately resulting in overgrowth by a pathogen. For this reason, we compared microbial community diversity and composition, growth rates, and metabolic features in infants that did and did not develop NEC. A clear finding of this study, and evident from prior research (17), is that microbial communities associated with infants that develop NEC are of lower diversity compared with control infants. However, this was due to the frequency of antibiotics administration for NEC infants. Regardless of the cause of the lower-diversity communities, microbial activities throughout the colonization process, including during periods of antibiotics administration, are likely important to infant health.

Several different species had higher relative abundance in infants that developed NEC, but none of these species were consistently associated with the disease. The correlation could be the consequence of the loss of other organisms from the community rather than their higher absolute abundance. Interestingly, *Veillonella spp.* were found to replicate more quickly in NEC versus control infants. This may be medically important, but additional examples are needed to establish a link between rapid growth and NEC.

Surprisingly, whether or not an infant developed NEC was not correlated with overall proteome composition. However, there were specific proteins that were associated with NEC, notably several involved in iron scavenging. Given that this is an important process often associated with pathogenesis, it is possible that increased activity of iron scavenging pathways could contribute to organism proliferation and onset of NEC. In addition, the *Klebsiella pneumoniae* proteome was correlated with NEC, including a protein involved in transport of iron. This is intriguing considering the prior finding that supplementation of lactoferrin, an abundant breast milk protein involved in modulating iron levels in the gut, decreases risk of developing necrotizing enterocolitis (22, 23). Overall, these findings indicate that fine-scale, species-specific proteins are important for understanding disease onset. Although the microbial community, and specific microbial proteins were correlated with NEC, no individual organism or protein was significantly more abundant in all cases. This finding supports the hypothesis that NEC is a multifaceted disease with multiple routes that lead to onset.

Although species-specific proteome profiles were correlated with community composition, they were largely infant specific. This is an interesting observation because it implies a feedback between human physiological conditions in the gut, which likely vary substantially from infant to infant and over time, and microbiome function.

## Methods

### Sample collection and metagenome sequencing

Samples were collected, processed for metagenome sequencing, and sequenced as part of three prior studies (**accession numbers in Supplemental File 1a**) (3, 4, 20). Stool samples were collected from infants and stored at −80°C. DNA was extracted from frozen fecal samples using the MO BIO PowerSoil DNA Isolation Kit, with modifications (4). DNA libraries were sequenced on an Illumina HiSeq for 100 or 150 cycles (Illumina, San Diego, CA). All samples were collected with parental consent.

### Metagenome assembly and genome binning

We re-assembled and analyzed metagenomes generated as part of a prior study, referred to as NIH1 (4). The data were processed in a manner consistent with the two other prior studies analyzed, referred to as NIH2 (20) and NIH3 (3). All raw sequencing reads were trimmed using Sickle (https://github.com/najoshi/sickle). Each metagenome was assembled separately using IDBA_UD (24). Open reading frames (ORFs) were predicted using Prodigal (25) with the option to run in metagenome mode. Predicted protein sequences were annotated based on USEARCH (–ublast) (26, 27) searches against UniProt (28), UniRef100 (29), and KEGG (30, 31). Scaffold coverage was calculated by mapping reads to the assembly using Bowtie2 (32) with default parameters for paired reads.

Scaffolds from NIH1 infants were binned to genomes using Emergent Self-Organizing Maps (ESOMs) generated based on time-series abundance profiles (15, 33). Reads from every sample were mapped independently to every assembly using SNAP (34), and the resulting coverage data were combined. Coverage was calculated over non-overlapping three Kbp windows. Coverage values were normalized first by sample, and then the values for each scaffold fragment were normalized from zero to one. Combining coverage data from scaffolds assembled from different samples prior to normalization made it possible to generate a single ESOM map for binning genomes assembled independently from each sample. ESOMs were trained for ten epochs using the Somoclu algorithm (35) with the option to initialize the codebook using Principal Component Analysis (PCA). Genomes were binned by manually selecting data points on the ESOM map using Databionics ESOM Tools (36). Binning was aided by coloring scaffold fragments on the map based on BLAST (37) hits to the genomes assembled in the prior study.

As part of the NIH2 and NIH3 studies, scaffolds were binned based on their GC content, DNA sequence coverage, and taxonomic affiliation using ggKbase tools (ggkbase.berkeley.edu). Genome bins from all three datasets were classified based on the consensus of taxonomic assignments for predicted protein sequences. Genome completeness and contamination were estimated for all genomes using CheckM with the taxonomy_wf option (38). Genomes with extra single copy genes, but with ≤175 fragments/Mbp (normalized for contamination) that were estimated to be ≥75% complete were manually curated based on scaffold GC content and coverage.

### Clustering genomes into sub-species groups

Genomes were clustered into sub-species groups based on sharing ≥98% average nucleotide identity (ANI), as estimated by MASH (39). Representative genomes were selected for each cluster as the largest genome with the highest expected completeness and smallest amount of contamination. Genomes were classified based on the lowest possible consensus of taxonomic assignments for predicted protein sequences.

Taxonomic assignments for representative genomes were checked manually based on hits to ribosomal protein S3, or visual inspection of protein taxonomic assignments. In order to identify cases in which the same bacterial strain was present in multiple samples, sub-species groups were further analyzed with the ANIm algorithm (40) implemented in dRep (41).

### Measuring microbial community abundance and replication rates

In order to achieve accurate abundance and replication rate measurements from read mapping, databases of representative genomes were created for each sample. Each database was constructed in order to include a representative genome from important sub-species groups. Priority was given to high-quality draft genome sequences reconstructed from the same sample. Genomes were classified as high-quality draft based on the requirements for iRep replication rate analysis (https://github.com/christophertbrown/iRep): ≥75% complete, ≤2.5% contamination, and ≤175 scaffolds per Mbp of sequence (21). Genomes were selected to represent sub-species groups using the following priority scheme: 1) high-quality draft genome assembled from the same sample, 2) high-quality draft genome from the same infant, 3) high-quality draft genome representative of sub-species group from any infant (if group had ≥5 representatives), 4) best genome from infant (if a genome was available). iRep was conducted using reads that mapped to genome sequences with ≤1 mismatch per read sequence. In cases where iRep values were ≥3, coverage plots were inspected and values were removed if there was evidence of strain variation.

We considered bacterial sub-species to be present in a sample if ≥97% of the genome was covered by an average of ≥2 reads. Abundance and iRep measurements were compared across samples by linking sample-specific representative genomes to sub-species groups. Relative abundance measurements for each sub-species group were calculated by converting DNA sequencing coverage values to a percentage. UniFrac (42) analysis was conducted based on rarefied abundance data and a tree constructed based on pairwise genome ANI values measured using MASH (-ms 5000000).

### Metaproteomics analysis

Metaproteomics sequencing was conducted on 0.3 g of stool as previously described (18). Each sample was suspended in 10 mL cold phosphate buffered saline. Samples were filtered through a 20 µm size filter to enrich for microbial cells and proteins. Microbial cells were collected by centrifugation, boiled in 4% sodium dodecyl sulfate for 5 minutes, and sonicated to lyse cells. The resulting protein extract was precipitated with 20% trichloroacetic acid at −80°C overnight. The protein pellet was washed with ice-cold acetone, solubilized in 8 M urea, reduced with 5 mM dithiothreitol, and cysteines were blocked with 20 mM iodoacetamide. Then sequencing grade trypsin was used to digest the proteins into peptides. Proteolyzed peptides were then salted and acidified by adjusting the sample to 200 mM NaCl, 0.1% formic acid, followed by filtering through a 10 kDa cutoff spin column filter to collect tryptic peptides.

Peptides were quantified by BCA assay and 50 µg peptides of each sample were analyzed via two-dimensional nanospray LC-MS/MS system on an LTQ-Orbitrap Elite mass spectrometer (Thermo Scientific). Each peptide mixture was loaded onto a biphasic back column containing both strong-cation exchange and reverse phase resins (C18). As previously described, loaded peptides were separated and analyzed using a 11-salt-pusle MudPIT protocol over a 22-h period (43). Mass spectra were acquired in a data-dependent mode with following parameters: full scans were acquired at 30 k resolution (1 microscan) in the Orbitrap, followed by CID fragmentation of the 20 most abundant ions (1 microscan). Charge state screening and monoisotopic precursor selection were enabled. Unassigned charge and charge state +1 were rejected. Dynamic exclusion was enabled with a mass exclusion width of 10 ppm and exclusion duration of 30 seconds. Two technical replicates were conducted for each sample.

Protein databases were generated for each infant from protein sequences predicted from assembled metagenomes. The database also included human protein sequences (NCBI Refseq_2011), common contaminants, and reverse protein sequences, which were used to control the false discovery rate (FDR). Collected MS/MS spectra were matched to peptides using MyriMatch v2.1 (44), filtered, and assembled into proteins using IDPicker v3.0 (45). All searches included the following peptide modifications: a static cysteine modification (+57.02 Da), an N-terminal dynamic carbamylation modification (+43.00 Da), and a dynamic oxidation modification (+15.99). A maximum 2% peptide spectrum match level FDR and a minimum of two distinct peptides per protein were applied to achieve confident peptide identifications (FDR <1%). To alleviate the ambiguity associated with shared peptides, proteins were clustered into protein groups by 100% identity for microbial proteins and 90% amino acid sequence identity for human proteins using USEARCH (26). Spectral counts were balanced between shared proteins, and proteins were considered to be present if ≥2 unique peptides were identified.

### Identification of putative protein families

Putative protein families were identified in order to track the presence and abundance of different protein types across samples. ORFs were first pre-clustered at 95% identity using USEARCH (-cluster_smallmem-target_cov 0.50 -query_cov 0.95 -id 0.95), and then all-versus-all protein searches were conducted (–ublast -evalue 10e-10 -strand both). Protein families were delineated from within the all-versus-all network graph using the MCL clustering algorithm (-I 2 -te 10) (46). The most common annotation observed across all protein sequences in the group was selected as the annotation for the putative protein family. Proteins were also grouped based on sharing 97% amino acid identity using USEARCH (-cluster_smallmem -target_cov 0.50 -query_cov 0.95 -id 0.97).

### Tracking human and bacterial protein abundances

Human and bacterial protein abundances were normalized using the weighted trimmed mean method from EdgeR (47). Species-specific proteomic profiles were normalized as the percent of total balanced spectral counts.

### Sample clustering and statistical analyses

Sample clustering was conducted based on microbial community abundance and iRep profiles, and bacterial protein family abundance profiles. In each case, the number of clusters was determined using the gap statistic (48), and then samples were grouped into the appropriate number of clusters using hierarchical clustering (average linkage method). Microbial community data was clustered based on weighted UniFrac distances, and protein data using Bray-Curtis distance. EdgeR was used to calculate statistically significant differences between conditions using quasi-likelihood linear modeling (glmQLFTest).

## Acknowledgements

Sample collection was approved by the University of Pittsburgh Institutional Review Board (PRO10090089). Funding was provided by National Institutes of Health grants R01-AI-092531 and R01-GM-103600.

## Author Contributions

MJM oversaw sample collection, RB collected all samples and managed metadata, and BF coordinated sample processing for DNA sequencing and proteomics analysis. CTB and MRO assembled and annotated the metagenome data. CTB and JFB carried out the genome binning and curation. CTB conducted the microbial community time series abundance and iRep analyses. WX and RLH generated the proteomics data, which was analyzed by CTB. CTB, MRO, and BCT provided bioinformatics support. CTB and JFB wrote the paper, and all authors provided input to the final text.

## Supplemental Materials

### Supplemental Figures

**Supplemental Figure 1 | Metagenome sequencing and metaproteomics conducted on microbiome samples collected from premature infants.** Frequency of samplecollection for metagenomics (**a**) and metaproteomics (**b**) based on infant day of life (DOL). **c**, Metagenome sequencing, and **d**, the percentage of each metagenome represented by assembled genome sequences ≥50% complete with ≤5% contamination. **e**, The number of proteomics spectral counts that could be uniquely assigned to human or bacterial proteins. **f**, The percent of predicted proteins that could be detected in each sample. **g**, The percent of species-specific proteomes that could be detected for species where ≥10% of the proteome could be detected in at least one sample. **h**, Histogram showing the distribution of the maximum percent of the proteome detected for all species present in each sample.

**Supplemental Figure 2 | ESOM genome binning.** Genome binning was conducted based on Emergent Self-Organizing Map (ESOM) clustering of scaffolds assembled from individual metagenomes. Data points represent 3 Kbp fragments of assembled scaffolds. Coloring is based on the species-level assignment of reconstructed genomes. The map is periodic, and red boxes indicate a single period.

**Supplemental Figure 3 | Infants that developed NEC and healthy controls are colonized by genetically similar bacteria.** Presence (dark boxes) and absence (white boxes) of members of bacterial sub-species in microbial communities from different infants. Sub-species were identified based on sharing ≥98% genome average nucleotide identity (ANI), and were determined to be present if ≥97% of the genome was covered by an average of ≥2 reads. Hierarchical clustering was conducted based on unweighted UniFrac distances calculated between infant genome inventories.

**Supplemental Figure 4 | Studied infant gut microbial communities associate into seven primary community types. a**, Hierarchical clustering was conducted based on the abundance of bacterial sub-species using weighted UniFrac distances. Microbial community types are identified by colored boxes. Metadata are shown for each sample, and indicated with an asterisk if significantly correlated with microbial community abundance data (PERMANOVA or Mantel test p-value ≤0.01). **b-i**, PCoA clustering of microbial communities with associated metadata: antibiotics administration (**b**), infant (**c**), developmental age (**d**; number of days since conception: gestational age + day of life, GA + DOL), proteome type (**e**), iRep type (**f**), infant health (**g**), days prior to NEC diagnosis (**h;** DOL – NEC diagnosis), and human proteome type (**i**).

**Supplemental Figure 5 | Microbial community abundance and replication rate profiles.** Relative abundance (bars) and iRep replication rate (scatter plot) values for bacterial sub-species colonizing studied premature infants. The five days following antibiotics administration are indicated with a color gradient.

**Supplemental Figure 6 | Microbial community types are distinguished by their abundant members.** Rank abundance curves showing the average and range (95% confidence interval) of relative abundance values for sub-species groups associated with each community type.

**Supplemental Figure 7 | Hierarchical clustering of genomes for members of the same sub-species group.** dRep results show ANI clustering of assembled genomes. Genome names indicate the metagenome that each genome was assembled from (**see Supplemental File 1b**). Clustering dendrograms show that most infants are colonized by different strains.

**Supplemental Figure 8 | Multiple species have infant-specific proteome profiles. a**, Analysis of *Veillonella spp.* genomes shows the presence of four different species. **b-e**, Proteome profiles for different species colonizing premature infants. Hierarchical clustering was conducted based on all detected protein families, and shows that strains colonizing different infants typically have distinct proteomic profiles. Infant and species metadata are shown for each sample. Metadata significantly correlated with the species proteome are indicated with an asterisk (PERMANOVA or Mantel test p-value ≤0.01). Protein families that correlated with at least one infant are shown in the heatmap (edgeR q-value ≤0.01). Samples colonized by the same strain are shown with colored text.

**Supplemental Figure 9 | Proteome types are distinguished by the abundance of proteins from different KEGG modules.** Hierarchical clustering of proteome types was conducted based on the abundance of proteins associated with KEGG modules. The relative abundance of proteins associated with each module was summed for each sample, and then the average was taken across all samples associated with each proteome type.

### Supplemental Files

**Supplemental File 1a | DNA sequencing and metaproteomics statistics. Supplemental File 1b | Genomes reconstructed from metagenomes.**

**Supplemental File 1c | Species iRep replication rates and statistical analysis. Supplemental File 1d | Species-specific microbial protein family abundance and statistical analysis.**

**Supplemental File 1e | Correlation of species-specific protein family abundances with iRep replication rates and gestational age corrected day of life (GA + DOL).**

**Supplemental File 1f | Species relative abundance and statistical analysis.**

**Supplemental File 1g | Microbial protein family abundance and statistical analysis.**

**Supplemental File 2 | Scaffolds binned to reconstructed genomes.**

**Supplemental File 3 | Proteins assigned to putative families.**

**Supplemental File 4 | Metaproteomics spectral counts.**

